# A Tetratricopeptide Repeat Protein Regulates Carotenoid Biosynthesis and Chromoplast Development in Monkeyflowers (*Mimulus*)

**DOI:** 10.1101/171249

**Authors:** Lauren E. Stanley, Baoqing Ding, Wei Sun, Fengjuan Mou, Connor Hill, Shilin Chen, Yao-Wu Yuan

## Abstract

The incredible diversity of floral color and pattern in nature is largely determined by the transcriptional regulation of anthocyanin and carotenoid biosynthetic genes. While the transcriptional control of anthocyanin biosynthesis is well understood, little is known about the factors regulating the carotenoid biosynthetic pathway in flowers. Here, we characterize the *Reduced Carotenoid Pigmentation 2* (*RCP2*) locus from two monkeyflower (*Mimulus*) species, the bumblebee-pollinated *M. lewisii* and hummingbird-pollinated *M. verbenaceus*. We show that loss-of-function mutations of *RCP2* cause drastic down-regulation of the entire carotenoid biosynthetic pathway in these species. Through bulk segregant analysis and transgenic experiments, we have identified the causal gene underlying *RCP2*, encoding a tetratricopeptide repeat (TPR) protein that is closely related to the *Arabidopsis* Reduced Chloroplast Coverage (REC) proteins. RCP2 appears to regulate carotenoid biosynthesis independently of RCP1, a previously identified R2R3-MYB master regulator of carotenoid biosynthesis. We show that RCP2 is required for chromoplast development and suggest that it most likely regulates the expression of carotenoid biosynthetic genes through chromoplast-to-nucleus retrograde signaling. Furthermore, we demonstrate that *M. verbenaceus* is just as amenable to chemical mutagenesis and *in planta* transformation as the more extensively studied *M. lewisii*, making these two species an excellent platform for comparative developmental genetics studies of two closely related species with dramatic phenotypic divergence.

## INTRODUCTION

Most flowers are colored by two classes of pigments: the red, pink, purple, or blue anthocyanins, and the yellow, orange, or red carotenoids. The hydrophilic anthocyanins are usually stored in the vacuoles of petal cells, whereas the hydrophobic carotenoids accumulate in chromoplasts as various lipoprotein structures (e.g., plastoglobules, crystals, fibrils). Frequently a plant can produce both pigment types in the same flower, forming contrasting spatial patterns that serve as nectar guides for animal pollinators (Glover, 2014). Common examples among horticultural plants include pansies, primroses, lantanas, and hibiscus, to name but a few. As an example in nature, the vast majority of the ∼160 species of monkeyflowers (*Mimulus*) (Barker et al., 2012) produce both anthocyanins and carotenoids in their petals with striking patterns. These observations indicate that many plant genomes contain a full set of functional genes encoding the anthocyanin and carotenoid biosynthetic pathways. The tremendous diversity of floral pigmentation pattern, then, is largely due to when and where these pathway genes are expressed. As such, identifying the transcriptional regulators of these pigment biosynthetic pathways are critically important to understanding the developmental mechanisms of pigment pattern formation and the molecular bases of flower color variation.

The transcriptional control of anthocyanin biosynthesis is well understood. A highly conserved MYB-bHLH-WD40 (MBW) protein complex has been shown to coordinately activate all or some of the anthocyanin biosynthetic pathway genes in multiple plant systems (Paz-Ares et al., 1987; Ludwig et al., 1989; Martin et al., 1991; Goodrich et al., 1992; de Vetten et al., 1997; Quattrocchio et al., 1998; Borevitz et al., 2000; Spelt et al., 2000; Schwinn *et al.*, 2006; reviewed in Davies *et al*., 2012; Glover, 2014). Among the three components, the R2R3-MYB often displays tissue-specific expression and causes spatial patterning of anthocyanin deposition in flower petals (Shang et al., 2011; Albert et al., 2011; Yuan et al., 2014; Martins et al., 2016). In contrast, little is known about the transcriptional regulators of the carotenoid biosynthetic pathway (CBP) (Ruiz-Sola & Rodríguez-Concepción, 2012; Yuan et al. 2015), particularly in flowers.

The best characterized transcriptional regulators of carotenoid biosynthesis are the phytochrome-interacting factors (PIFs) in *Arabidopsis* (Toledo-Ortiz et al., 2010). PIFs directly bind the promoter of the *phytotene synthase* (*PSY*) gene and repress its expression in dark-grown seedlings. During deetiolation, light-triggered degradation of PIFs leads to rapid derepression of *PSY* and massive production of carotenoids in the greening seedlings. However, the significance of PIFs in regulating carotenoid pigmentation in flowers is unclear. In fact, there are good reasons to suspect that PIFs do not play an important role in flower pigmentation: several studies have shown that carotenoid production during petal development involves coordinated activation of multiple CBP genes (Giuliano et al., 1993; Moehs *et al*., 2001; Zhu et al., 2002; Chiou *et al*., 2010; Yamagishi *et al*., 2010; Yamamizo *et al*., 2010), whereas PIFs regulate only *PSY* but none of the other CPB genes in *Arabidopsis* (Toledo-Ortiz et al., 2010).

To identify transcriptional regulators of floral carotenoid pigmentation, we employed a new genetic model system, the monkeyflower species *Mimulus lewisii*. The ventral (lower) petal of *M. lewisii* flowers has two yellow ridges that are pigmented by carotenoids (Figure 1A and 1B), serving as nectar guides for bumblebee pollinators (Owen and Bradshaw, 2011). We carried out an ethyl-methanesulfonate (EMS) mutant screen using the inbred line LF10 for reduced carotenoid pigmentation in the nectar guides, and recovered several recessive mutants with coordinated down-regulation of the entire CBP compared to the wild-type (Sagawa et al., 2016). Complementation crosses suggested that these mutants represent two different loci, *RCP1* (*Reduced Carotenoid Pigmentation 1*) and *RCP2*. *RCP1* encodes a subgroup-21 R2R3-MYB that is clearly distinguishable from the anthocyanin-activating R2R3-MYBs (subgroup-6) and is specifically expressed in the yellow nectar guides (Sagawa et al., 2016) in *M. lewisii*.

**Figure 1.**
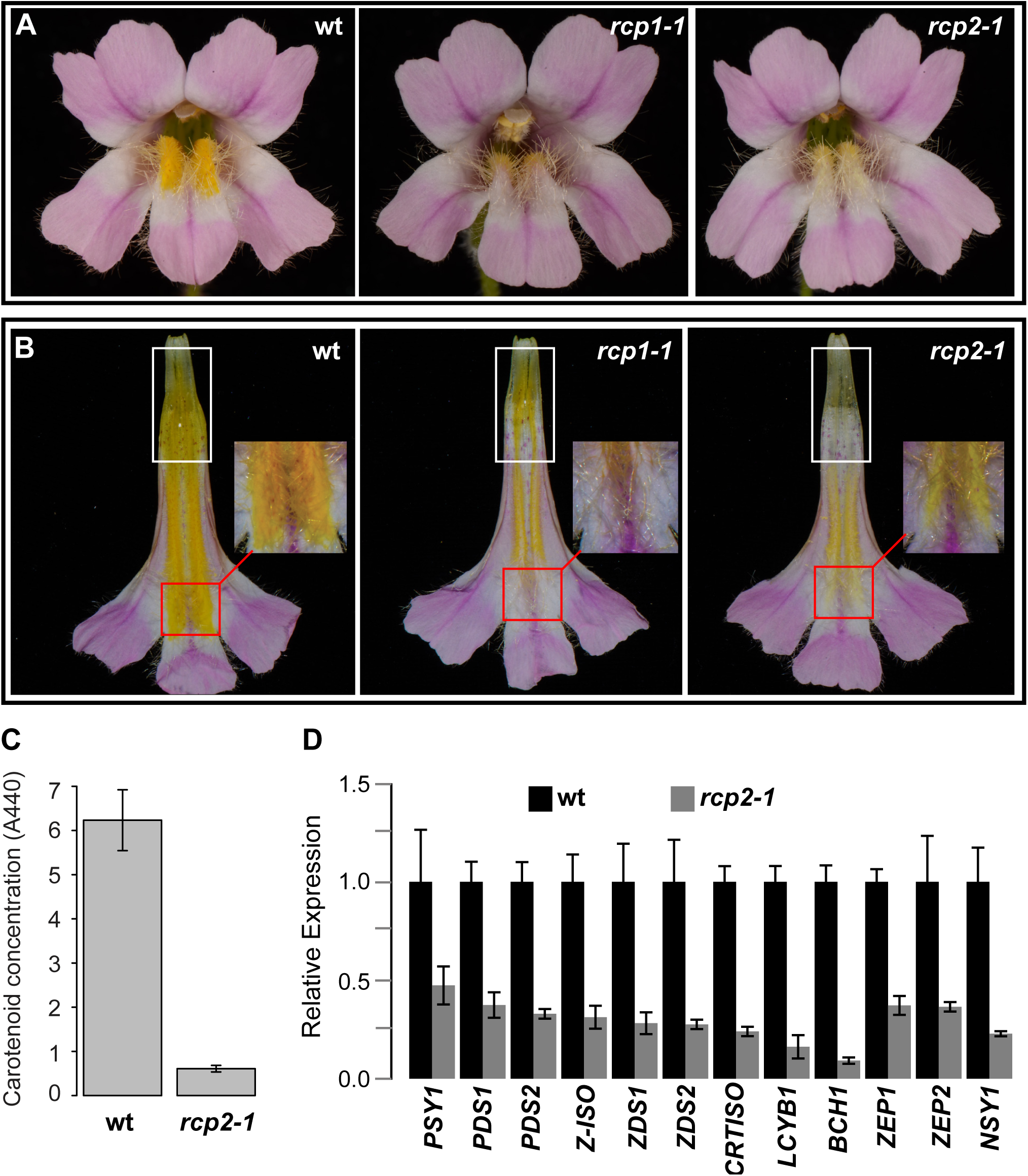
Reduced carotenoid pigmentation phenotypes in *Mimulus lewisii*. (**A**) Front view of the wild-type (wt), *rcp1-1*, and *rcp2-1* flowers. (**B**) Nectar guide view. White and red boxes indicate the base and throat of the corolla tube, respectively. (**C**) Carotenoid concentration, as estimated by absorbance measurements at 440 nm. Error bars are 1 SD (n = 8). (**D**) Relative transcript level of the CBP genes in *rcp2-1* compared to wt, as determined by qRT-PCR. Error bars are 1 SD (n = 3).

The primary goal of this study is to characterize the *RCP2* locus. Through bulk segregant analysis and transgenic experiments, we have identified the causal gene of *RCP2*, encoding a tetratricopeptide repeat (TPR) protein homologous to the Reduced Chloroplast Coverage (REC) proteins in *Arabidopsis*. Loss-of-function *REC* mutants have reduced chlorophyll content and smaller chloroplast compartment size compared to wild-type (Larkin et al., 2016). Our analyses show that *RCP2* is required for chromoplast development and carotenoid biosynthesis in the flowers of both *M. lewisii* and a close relative, *M. verbenaceus*. We suggest that the coordinated down-regulation of the nuclear-encoded CBP genes in the loss-of-function *rcp2* mutant likely results from chromoplast-to-nucleus retrograde signaling, which appears to be independent of *RCP1* function.

## RESULTS

### *rcp2-1* displays a distinct and stronger phenotype than *rcp1-1*

We recovered three independent *rcp2* alleles from the previous mutant screen. *rcp2-1* and *rcp2-2* are indistinguishable phenotypically, whereas *rcp2-3* displays a slightly weaker phenotype (Supplemental Figure 1). Like the *rcp1-1* mutant, *rcp2-1* has reduced carotenoid content in the nectar guides compared to the wild-type (Figures 1A and 1B). However, *rcp2-1* can be readily distinguished from *rcp1-1* in two aspects. First, the total carotenoid content in the nectar guides of *rcp2-1* is ∼10-fold lower than the wild-type (Figure 1C), whereas *rcp1-1* is only ∼4.4-fold lower (Sagawa et al., 2016). Second, the residual carotenoid pigments in *rcp1-1* and *rcp2-1* show distinct spatial distributions. At the base of the corolla tube (Figure 1B, white boxes), carotenoid pigments are completely lacking in *rcp2-1* but present in *rcp1-1*. In contrast, at the throat of the corolla tube (Figure 1B, red boxes), carotenoid pigments are completely lacking in *rcp1-1* but present at low concentration in *rcp2-1*, giving a cream color. These spatial distributions of residual pigments are consistent among allelic mutants within each complementation group (Supplemental Figure 1).

To test whether RCP2 regulates CBP genes at the transcriptional level, we performed qRT-PCR experiments on nectar guide tissue at the 15-mm corolla developmental stage (the stage at which the CBP genes have their highest expression; this is the same stage used previously for RCP1). Compared to wild-type, the *rcp2-1* mutant showed a coordinated down-regulation of the entire CBP (Figure 1D), with a 3- to 4-fold decrease in expression of most CBP genes and ∼10-fold decrease in *BCH1* expression. These results suggest that RCP2 is involved in the transcriptional regulation of CBP genes in the nectar guides. Consistent with the more severe reduction in total carotenoid content, the extent of CBP gene down-regulation is stronger in *rcp2-1* than in *rcp1-1* (Sagawa et al., 2016).

### *RCP2* encodes a TPR protein

To identify *RCP2*, we performed bulk segregant analysis by Illumina sequencing of an F2 population, which was derived from a cross between *rcp2-1* (in the LF10 background) and the mapping line SL9 (see METHODS). A conspicuous peak was detected on pseudoscaffold 11 (Figure 2A), corresponding to a 70-Kb region in the LF10 genome (LF10g_v1.8 scaffold 278: 1,215,000 – 1,285,000). Remapping the Illumina reads to this 70-Kb interval of the LF10 genome revealed only one mutation in the entire region. This mutation causes a premature stop codon at the beginning of the second exon of a gene encoding a TPR protein of 1,794 amino acids (Figure 2B). Sequencing the independent *rcp2-2* and *rcp2-3* alleles showed that both contain nonsynonymous amino acid replacements at highly conserved sites (Figure 2B and Supplemental Figure 2), corroborating the candidacy of this gene.

**Figure 2.**
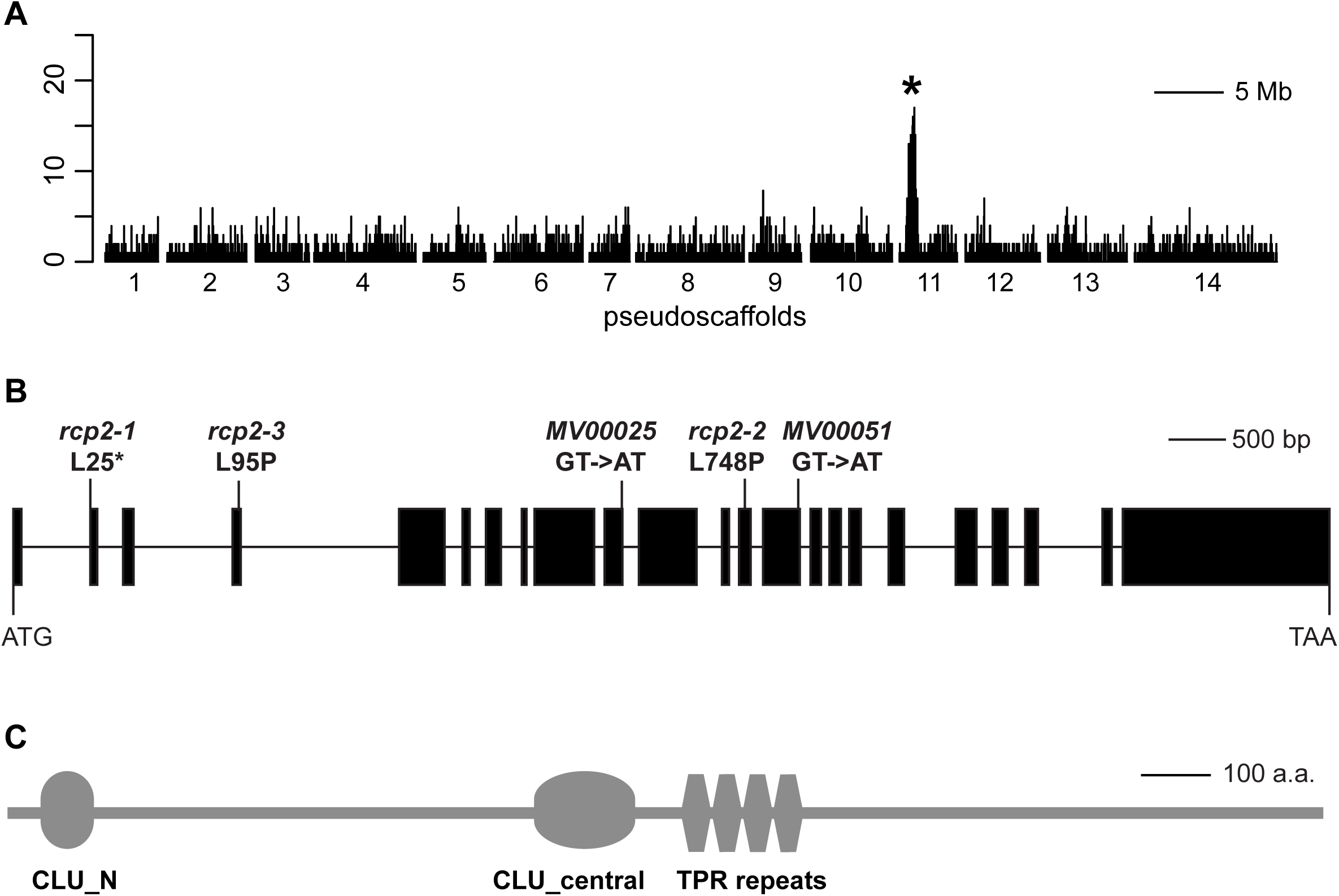
Identification of *RCP2*. (**A**) Whole genome scan for regions enriched in homozygous SNPs. (**B**) Schematic *RCP2* gene map highlighting mutations from *M. lewisii (rcp2-1*, *rcp2-2*, *and rcp2-3)* and *M. verbenaceus* (*MV00025* and *MV00051*). Black boxes: exons; lines: introns. (**C**) RCP2 protein schematic showing the key domains.

To further verify that this TPR gene is *RCP2*, we wanted to perform two transgenic experiments: (1) transform a genomic copy of the wild-type LF10 allele into the *rcp2-1* mutant background to restore the wild-type phenotype; and (2) transform an RCP2 RNAi construct into the wild-type LF10 background to recapitulate the mutant phenotype. Unfortunately, the extremely large size of the genomic fragment (∼11.5 kb excluding 5’ and 3’ regulatory sequences) prevented us from performing the rescue experiment. Therefore, we proceeded with the RNAi experiment only. We constructed an RNAi plasmid using a 408-bp fragment in the last exon, which has a unique nucleotide sequence as determined by BLASTing against the LF10 genome assembly, and transformed it into wild-type LF10. We obtained 127 stable transgenic plants, 96% of which closely resemble the *rcp2-1* mutant, including the complete lack of yellow pigments at the base of the corolla tube and the creamy yellow color at the throat of the corolla tube (Figure 3A). Evaluation of the expression level of this *TPR* gene in three of the RNAi lines showed ∼95% knockdown (Figure 3C), and qRT-PCR confirms that all of the CBP genes are down-regulated in the RNAi lines to a similar degree as in the *rcp2-1* mutant (Figure 3B). Taken together, these results suggest that this *TPR* gene is indeed *RCP2* and *rcp2-1* is likely to be a null allele, as the translated protein would contain only 24 of the 1,794 amino acids (Figure 2B).

**Figure 3.**
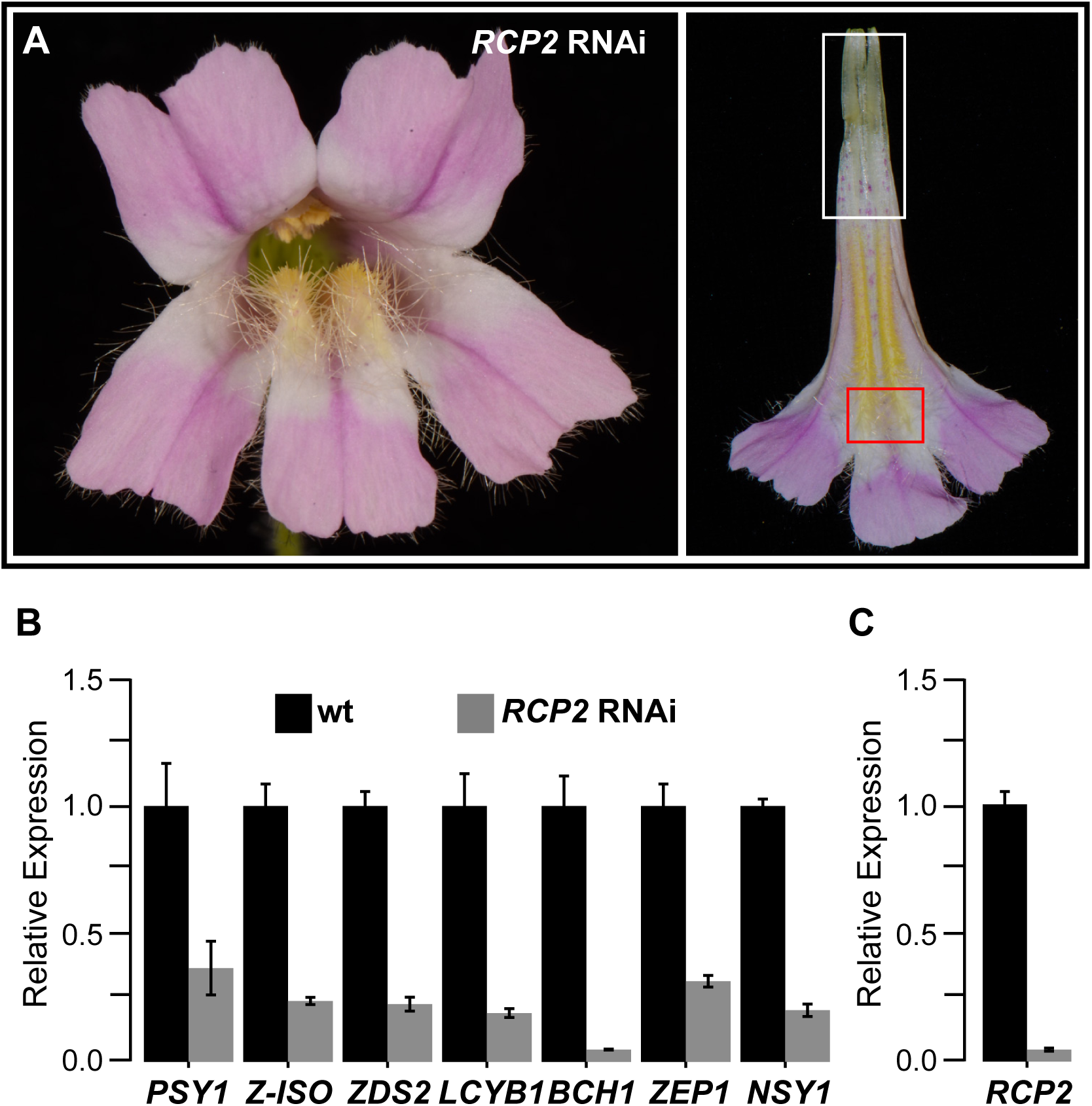
Characterization of *M. lewisii RCP2* RNAi lines. (**A**) Front view and nectar guide view of a representative *RCP2* RNAi line. (**B**) Relative transcript level of a subset of CBP genes in three *RCP2* RNAi lines compared to wild-type (wt), as determined by qRT-PCR. (C) qRT-PCR measurement of the *RCP2* expression level in three *RCP2* RNAi lines. Error bars are 1 SD (n = 3).

Blasting the RCP2 protein against the NCBI Conserved Domain Database (Marchler-Bauer *et al*., 2015) revealed three conserved domains –– a TPR domain with four TPR repeats, a <u>CLU</u>stered mitochondria protein N-terminal domain (CLU_N), and a CLU central domain (CLU_central) (Figure 2C) –– but no recognizable DNA-binding domain. TPR repeats are found in a wide range of proteins as scaffolds mediating protein-protein interactions (D’Andrea & Regan, 2003; Zeytuni and Zarivach, 2012; Bohne et al., 2016). In contrast, no specific functions have been assigned to the CLU domains to date. There are two closely related paralogs of *RCP2* in the *M. lewisii* genome, *RCP2-L1* and *RCP2-L2* (Figure 4A and Supplemental Figure 2). Phylogenetic analysis indicates that the divergence of these three paralogs predates the split between monocots and eudicots: there is a corresponding ortholog for each of the three genes in *Arabidopsis* and *Brachypodium* (Figure 4A). The *Arabidopsis* homologs (*REC1*: *AT1G01320*; *REC2*: *AT4G28080*; *REC3*: *AT1G15290*) have been shown to help establish the size of the chloroplast compartment in *Arabidopsis* leaf cells (Larkin et al., 2016). *RCP2* is most closely related to *REC2* (Figure 4A).

**Figure 4.**
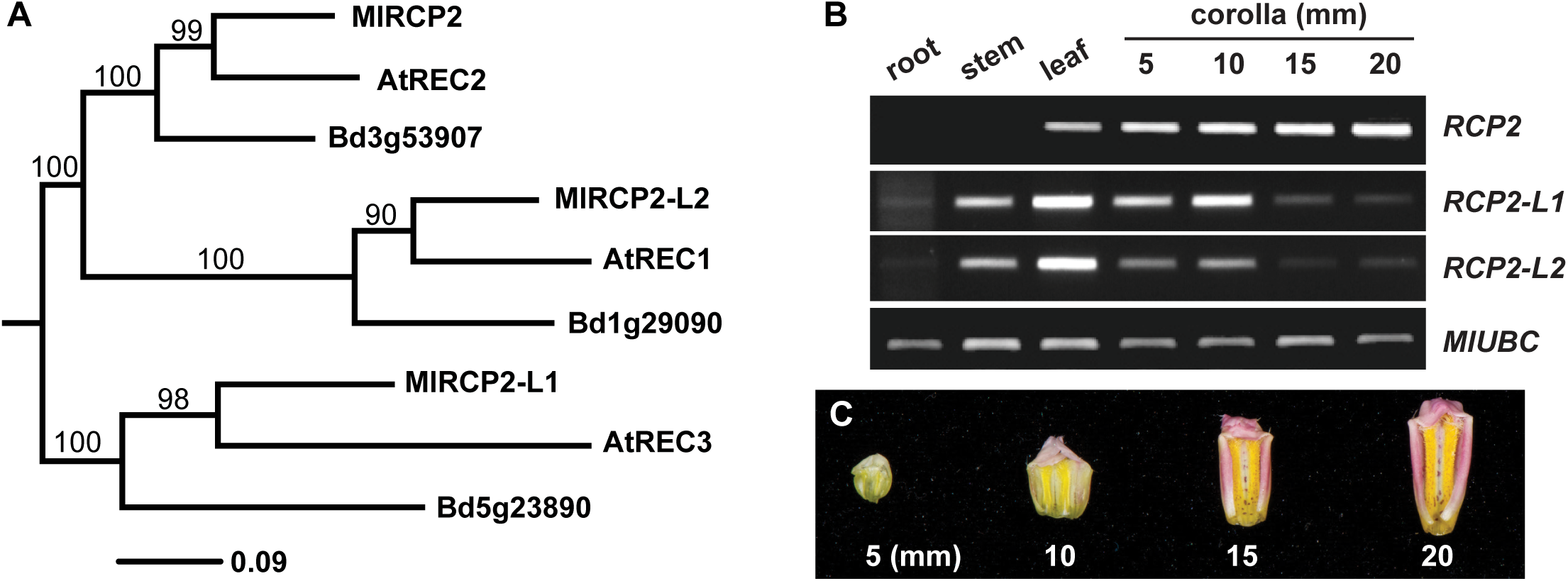
Divergence of *RCP2* and *RCP2-like* genes. (**A**) A maximum likelihood (ML) phylogeny of RCP2 and related proteins in *Mimulus lewisii*, *Arabidopsis thaliana*, and *Brachypodium distachyon*. ML analysis was conducted using the RAxML web-server (http://embnet.vital-it.ch/raxml-bb/), with the JTT amino acid substitution matrix and the GAMMA model of rate heterogeneity. Clade support was estimated by 100 bootstrap replicates. The tree is rooted by midpoint rooting. (**B**) Qualitative RT-PCR (28 cycles) of *RCP2* and *RCP2-like* genes in various tissues and corolla developmental stages of the wild-type *M. lewisii. MlUBC* was used as the reference gene. (**C**) Carotenoid accumulation in the nectar guides of wild-type *M. lewisii*.

The existence of three closely related paralogs with presumably redundant functions raised the question as to why mutations in a single gene, *RCP2*, result in such a strong phenotype in floral carotenoid pigmentation. We hypothesized that *RCP2* and the *RCP2*-like genes have evolved different expression patterns. To test this, we performed RT-PCR at different stages of corolla development and in different tissues (Figure 4B). *RCP2* is primarily expressed in the corolla and its expression gets progressively stronger as the corolla grows larger and the yellow color in the nectar guides becomes more intense (Figure 4B and Figure 4C). In contrast, *RCP2-L1* and *RCP2-L2* are only expressed in the early stages of corolla development before the yellow color becomes conspicuous, suggesting a relatively minor role of these genes in floral carotenoid pigmentation compared to *RCP2*. On the other hand, these two genes are strongly expressed in the leaf, where *RCP2* is expressed relatively weakly, explaining the lack of obvious phenotype in the vegetative tissue of *rcp2-1*.

### *RCP2* regulates carotenoid biosynthesis independently of *RCP1*

The fact that *rcp2-1* and *rcp1-1* are similar both in their visible phenotypes and their effects on transcriptional regulation of the CBP genes led to our initial hypothesis that *RCP1* and *RCP2* are part of the same genetic pathway. Under this scenario, there are three possible relationships between *RCP1* and *RCP2*: (**i**) *RCP1* is upstream of *RCP2* (e.g., *RCP1* regulates *RCP2* expression); (**ii**) *RCP2* is upstream of *RCP1* (e.g., *RCP2* regulates *RCP1* expression); and (**iii**) The two genes act at the same regulatory level (e.g., RCP1 and RCP2 form a protein complex). We considered the third possibility most likely, as RCP2 contains a TPR domain known to promote protein-protein interaction and the formation of protein complexes.

To test the first two possibilities, we performed qRT-PCR to assay the transcript level of *RCP2* in the *rcp1-1* mutant, and transcript level of *RCP1* in *rcp2-1*, relative to the wild-type. We found no significant differences in these comparisons (Figure 5A), which suggests that *RCP1* and *RCP2* do not regulate each other’s transcription. This is in line with the very different expression patterns of the two genes: while *RCP1* expression is restricted to the nectar guides (Sagawa et al., 2016), *RCP2* is expressed in both nectar guides and petal lobes, with slightly higher levels in petal lobes than nectar guides (Supplemental Figure 3A). To test the third possibility, we performed a yeast-two hybrid assay between RCP1 and RCP2. When the bait plasmid pGBKT7-RCP2 (expressing the GAL4 DNA-binding domain) and the prey plasmid pGADT7-RCP1 (expressing the GAL4 activation domain) were brought together in yeast cells, they were unable to activate the reporter genes (Supplemental Figure 3B), indicating that the RCP1 and RCP2 proteins do not directly interact in yeast cells. This lack of protein interaction is also in line with their sub-cellular localization: transient expression assay in tobacco leaves showed that RCP1 is located exclusively in the nucleus (Figure 5B), whereas RCP2 is located in both the nucleus and the cytosol, consistent with the previously reported localization of *Arabidopsis* REC1 (Larkin et al., 2016).

**Figure 5.**
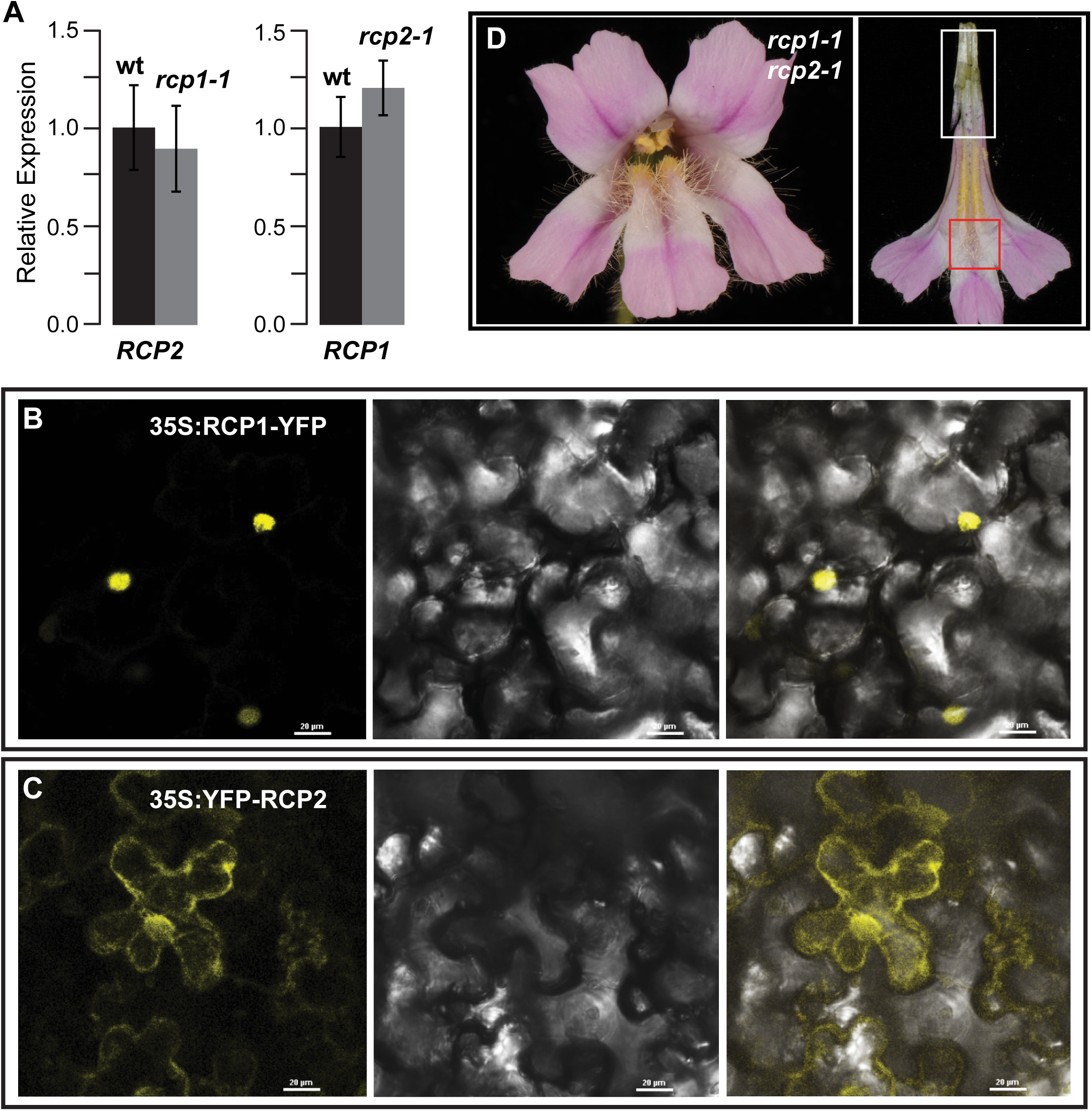
Lack of genetic interaction between *RCP1* and *RCP2*. (**A**) qRT-PCR experiments. Error bars are 1 SD (n = 3). (**B** and **C**) Sub-cellular localization of RCP1 and RCP2 proteins in *Nicotiana benthamiana* leaves. Left: green channel; middle: transmitted light; right: merged. (**D**) Front view and nectar guide view of the *rcp1-1 rcp2-1* double mutant.

In summary, none of these results support our initial hypothesis that *RCP1* and *RCP2* operate in the same genetic pathway. In other words, *RCP1* and *RCP2* most likely represent two independent pathways that regulate the transcript level of CBP genes. This inference is also supported by double mutant analysis: the *rcp1-1 rcp2-1* double mutant shows an additive phenotype regarding the spatial distribution of the residual carotenoids, which are completely absent at both the petal lobe base (as in *rcp2-1*) and the throat (as in *rcp1-1*) of the corolla tube (Figure 5D).

### *RCP2* is required for chromoplast development

If *RCP2* does not interact with *RCP1* in the same genetic pathway, then what is the function of *RCP2* in the regulation of carotenoid biosynthesis? The *RCP2* homologs in *Arabidopsis*, *REC1-REC3*, are involved in controlling the chloroplast compartment size (Larkin et al., 2016), and another *TPR* gene that is sister to the *REC* clade, *FRIENDLY* (Larkin et al., 2016), has been shown to control mitochondrial morphology and intracellular distribution (Logan et al., 2003; El Zawily et al., 2014). These observations prompted us to speculate that perhaps in the *M. lewisii* nectar guides, *RCP2* is involved in the development of another organelle, the chromoplast. This supposition seemed promising because chromoplasts and chloroplasts are known to be interconvertable (Egea et al., 2010; Li and Yuan, 2013), and chromoplast development is known to play an important role in carotenoid accumulation, although not necessarily in the transcriptional regulation of the CBP genes (Mustilli et al., 1999; Liu et al., 2004; Lopez et al., 2008; Galpaz et al., 2008).

To test this possibility, we examined the ultra-structure of the nectar guide epidermal cells by transmission electronic microscopy (TEM). In the wild-type, numerous rounded chromoplasts are pushed to the periphery of the cell by the vacuole (Figure 6A), and each chromoplast contains several electron-dense plastoglobuli, the main carotenoid-sequestering structures (Figure 6B). In sharp contrast, in the *rcp2-2* mutant, the chromoplasts on the cell periphery are skinny, irregularly shaped, and usually contain no plastoglobuli (Figure 6C). These results suggest that *RCP2* is required for proper chromoplast differentiation. However, these results alone do not distinguish between abnormal chromoplast differentiation and decreased carotenoid biosynthesis; which is the cause and which is the consequence? We reasoned that if decreased carotenoid biosynthesis is the cause, the *rcp1-1* mutant should also exhibit abnormal chromoplast development. However, the *rcp1-1* chromoplasts appear normal in shape, although the plastoglobuli are smaller in size and less electron-dense than in the wild-type (Figure 6D), as one would expect if carotenoids are less abundant. Based on these results, we conclude that in the *rcp2-2* mutant, abnormal chromoplast differentiation is most likely the cause and decreased carotenoid biosynthesis, through down-regulation of the CBP structural genes, is the consequence.

**Figure 6.**
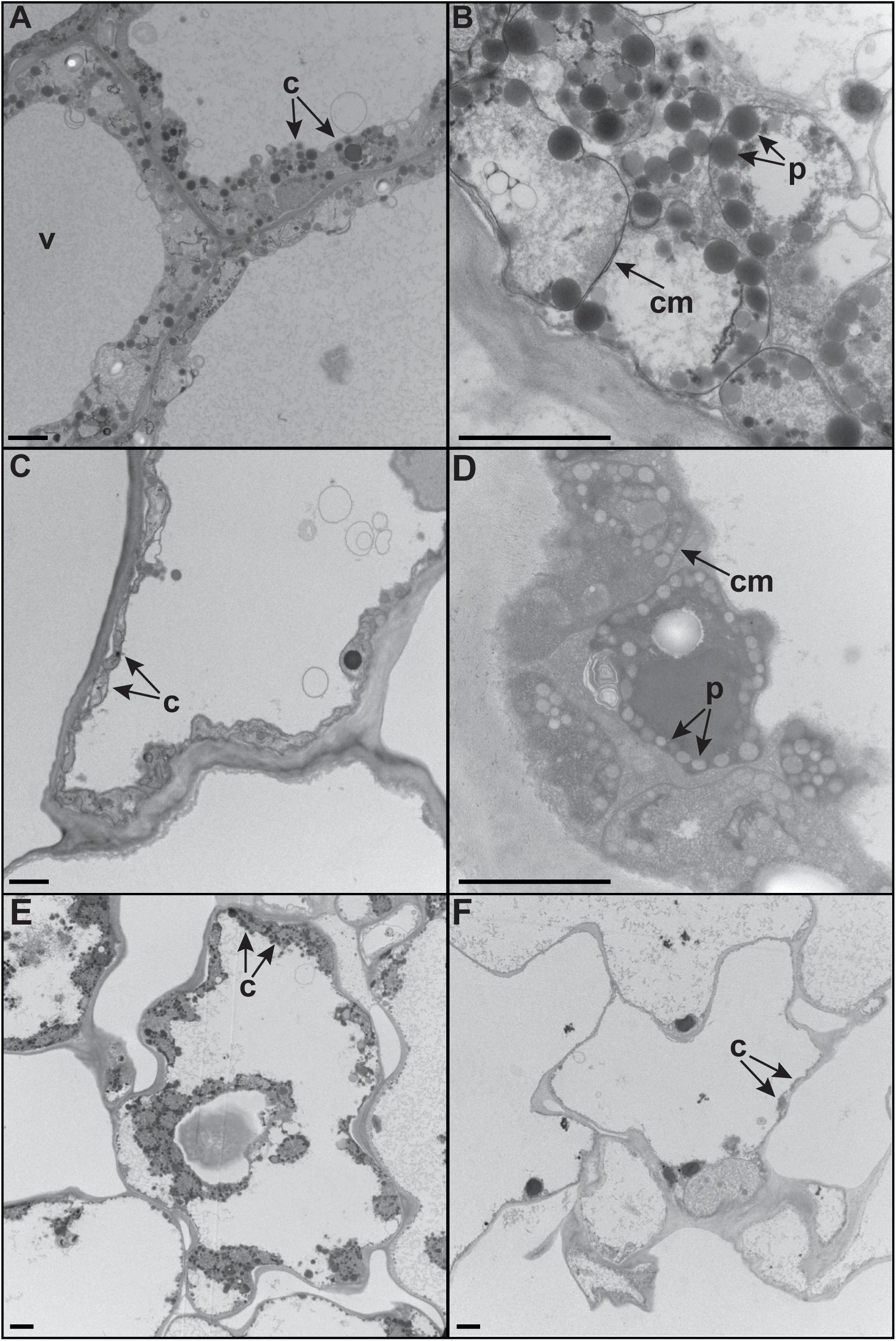
Transmission electron micrographs of chromoplasts. (**A**) Whole-cell view of the wild-type *M. lewisii* nectar guide upper epidermal cells. (**B**) Detailed view of individual chromoplasts of wild-type *M. lewisii*. (**C**) Whole-cell view of *rcp2-2*. (**D**) Detailed view of individual chromoplasts of *rcp1-1*. (**E** and **F**) Whole-cell view of the petal upper epidermal cells of wild-type *M. verbenaceus* (E) and *MV00051* (F). v = vacuole, c = chromoplast, p = plastoglobule, cm = chromoplast membrane. Scale bar is 2 μm.

### *RCP2* function is conserved in *M. verbenaceus*

Although *RCP2* is expressed in both the nectar guides and petal lobes, the phenotypic consequences of *RCP2* mutation in the petal lobes could not be revealed in *M. lewisii*, as a dominant repressor, *YELLOW UPPER* (*YUP*), is known to prevent carotenoid accumulation in *M. lewisii* petal lobes (Hiesey *et al*., 1971). To test whether the role of *RCP2* in carotenoid pigmentation is restricted to nectar guides or is more ubiquitous throughout the flower, we investigated *M. verbenaceus*, a hummingbird-pollinated species that is closely related to *M. lewisii* (Beardsley et al., 2003). *M. verbenaceus* is homozygous for the recessive *yup* allele, and accumulates carotenoids in both the petal lobe and nectar guide. The bright red color of the corolla results from the combination of high concentrations of carotenoids and anthocyanins (Figure 7A).

**Figure 7.**
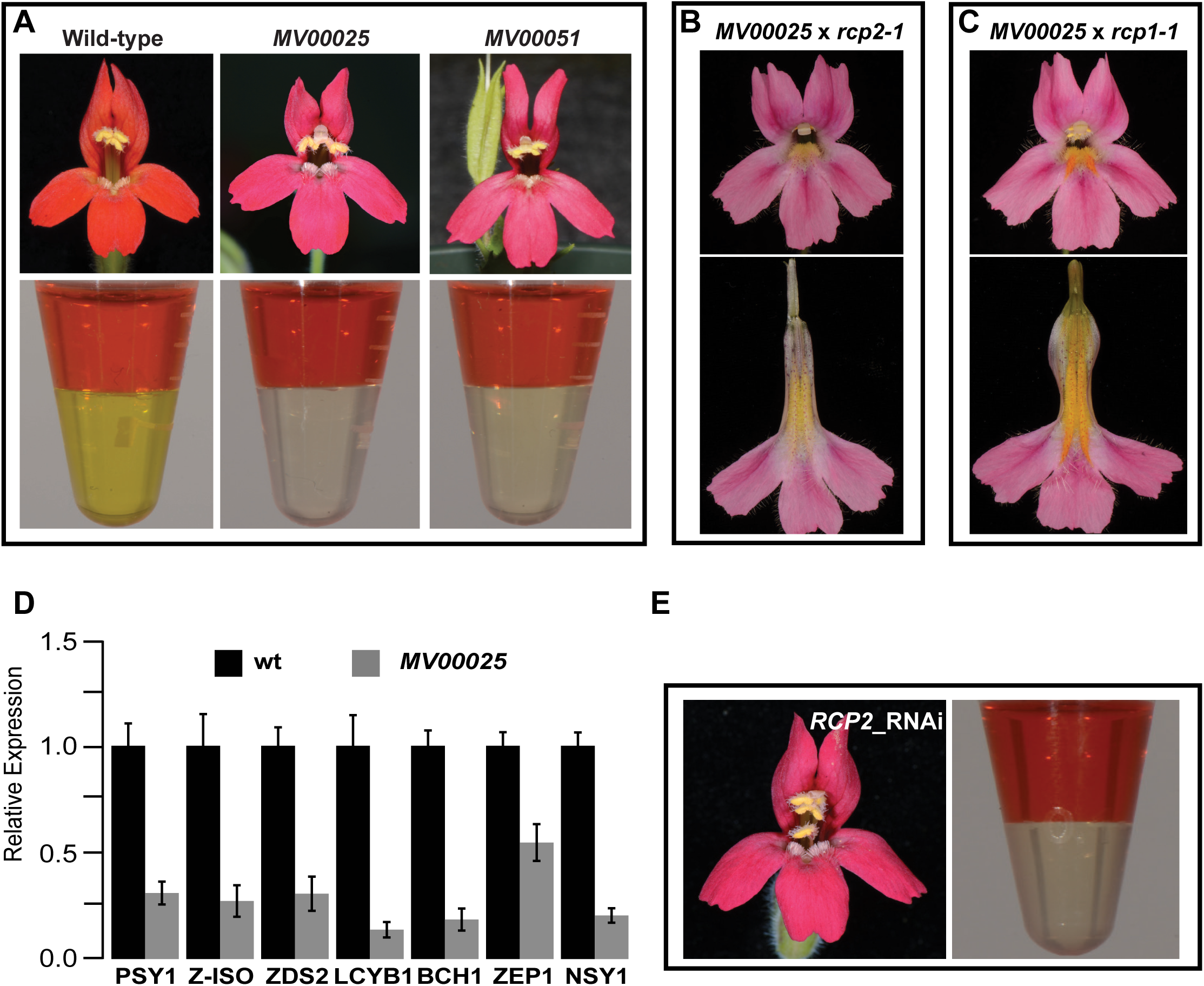
*RCP2* function in *Mimulus verbenaceus.* (**A**) Wild-type *M. verbenaceus* and the two *rcp2* mutants. Top row: front view of the flower. Bottom row: separation of anthocyanins (upper layer) and carotenoids (lower layer). (**B** and **C**) Complementation crosses. (**D**) qRT-PCR of a subset of the CBP genes in wild-type *M. verbenaceus* and the *rcp2* mutant *MV00025*. (E) Front view (left) and pigment separation (right) of a representative *RCP2* RNAi flower in *M. verbenaceus*.

Because we were interested in developing *M. verbenaceus* as a parallel model to *M. lewisii* for comparative developmental genetics studies, we performed a pilot EMS mutagenesis experiment using the *M. verbenaceus* inbred line MvBL. We screened 460 M1 families (∼20 M2 plants per family) and recovered more than 100 floral mutants, including two recessive mutants (*MV00025* and *MV00051*) that lack carotenoids in the entire corolla (Figure 7A; the magenta color is due to anthocyanins alone). Crossing these two mutants with each other and with *rcp2-1* produces flowers with very little carotenoid pigmentation in the nectar guides (Figure 7B). In contrast, crosses between the *M. verbenaceus* mutants and *rcp1-1* produce intense yellow color in the nectar guides (Figure 7C). These complementation crosses suggest that *MV00025* and *MV00051* are two additional *rcp2* alleles. The petal lobes of these F1 hybrids are without carotenoids (and therefore pink) regardless of the *RCP2* genotype, as they are heterozygous for the dominant *YUP* allele. Sequencing the *RCP2* coding DNA of the two *M. verbenaceus* mutants revealed independent intron splicing mutations (Figure 2B), corroborating the allelic test. Furthermore, the CBP genes are coordinately down-regulated in *MV00025* compared to the wild-type (Figure 7D), the chromoplasts of *MV00051* show similar abnormal phenotypes to the *rcp2-2* mutant (Figure 6E and F), and 37 of the 40 independent *RCP2* RNAi lines in *M. verbenaceus* recapitulated the mutant phenotype (Figure 7E). Taking all these results together, we conclude that the role of *RCP2* in regulating carotenoid biosynthesis and the expression of CBP structural genes is conserved between the two *Mimulus* species, and that *RCP2* function is ubiquitous throughout the corolla.

## DISCUSSION

In this study, we characterized the *RCP2* locus of *Mimulus lewisii* and its close relative *M. verbenaceus*. Based on genetic mapping, multiple independent mutant alleles, and RNAi knockdown experiments, we conclude that *RCP2* encode a TPR protein required for chromoplast development and CBP gene expression in *Mimulus*.

The RCP2 protein does not contain any known DNA-binding or activation domains. How, then, does it affect the transcript level of the CBP genes? The finding that mutations in *RCP2* disrupt chromoplast development (Figure 6) is interesting, as abnormal development of chloroplast is known to cause transcriptional down-regulation of photosynthesis-associated nuclear genes, an extensively studied process referred to as retrograde signaling (Susek et al., 1993; Mochizuki et al., 2001; Larkin et al., 2003; Strand et al., 2003; Sun et al., 2011; Xiao et al., 2012; reviewed in Leister, 2012; Chi et al. 2013; Chan et al., 2016). Given that chromoplasts and chloroplasts are interconvertible plastids (Egea et al., 2010; Li and Yuan, 2013), it is plausible there is also retrograde signaling from chromoplasts to nucleus affecting the transcription of carotenoid biosynthesis and accumulation genes.

Compared to chloroplast development and retrograde signaling, little is known about the nuclear regulation of chromoplast development and the communication between chromoplasts and the nucleus. The nuclear gene *Or* is known to be involved in converting colorless proplastids or leucoplasts into chromoplasts (Lu et al., 2006; Tzuri et al. 2015). The nuclear-encoded heat shock protein HSP21 has also been shown to be necessary for the chloroplast-to-chromoplast transition in ripening tomatoes (Neta-Sharir et al. 2005). However, no nuclear-encoded regulatory genes have been directly implicated in chromoplast-to-nucleus signaling. A previous proteomic analysis found that chromoplasts of tomato fruits are enriched in several components of the calcium signaling pathway, which is known to be involved in chloroplast retrograde signaling (Barsan et al. 2010). This suggests that perhaps chromoplast retrograde signaling utilizes some of the same pathways as chloroplast retrograde signaling. It would be interesting to transgenically manipulate components of these pathways in the *rcp2* mutant background.

Despite the clear importance of RCP2 and the homologous REC and FRIENDLY proteins in organelle development, morphology, and intra-cellular distribution (Logan et al., 2003; El Zawily et al., 2014; Larkin et al., 2016), the biochemical function of these proteins remains elusive. One way to approach this problem is to use the TPR domain as a bait to screen for interacting partners, which may provide new insights into the function of these putative protein complexes.

Another implication of this study is that floral CBP gene expression appears to be regulated by multiple pathways that ultimately converge. Though RCP1 and RCP2 do not operate in the same genetic pathway (Figure 5), both are required for the coordinated activation of the entire CBP, with the strongest effect on *BCH1* (Figure 1D; Sagawa et al., 2016). This type of redundant regulatory mechanism is certainly not unprecedented in plants. Flowering time, for example, is controlled by multiple distinct pathways (e.g., photoperiod, vernalization, autonomous, and gibberellic acid) that all converge on the floral integrators such as *FLOWERING LOCUS T* (Glover, 2014). It makes sense that plants have evolved multiple signaling pathways to regulate flowering time, since complex environmental cues must be sensed and integrated into an ultimate response –– flowering, which is essential for plant fitness. Why, then, would plants evolve multiple regulatory pathways to control a “nonessential” trait like floral carotenoid pigmentation? Many flowers, after all, do not produce carotenoids in their petals. It is important to note that in addition to coloring flowers and fruits, carotenoids play an indispensable role in photosynthesis and photoprotection. Therefore, leaf carotenoid composition is remarkably conserved across higher plants (Goodwin and Britton, 1988). Plants may have evolved multiple regulatory pathways to establish and maintain this optimized carotenoid composition in vegetative tissues in response to a variable environment; it was only later during plant evolution that these pathways were co-opted for flower coloration.

Another layer of redundancy exists at the gene level. Given the amino acid sequence similarity between RCP2 and its paralogs RCP1-L1 and RCP2-L2 (Supplemental Figure 2), it is reasonable to assume that they are at least partially functionally redundant. This redundancy may ensure proper carotenoid accumulation in vegetative tissues for photosynthesis, while allowing for innovation in non-green tissues like flowers. The evolution of tissue-specific expression patterns (i.e., *RCP2* in flowers and the other two paralogs in vegetative tissues; Figure 4B) may underlie this lability. *RCP2* and the *RCP2*-like genes therefore have great potential in generating natural variation in floral carotenoid pigmentation.

Finally, this study demonstrates that the hummingbird-pollinated *M. verbenaceus* is just as amenable to chemical mutagenesis and *in planta* transformation (Figure 7) as the more extensively studied, bumblebee-pollinated *M. lewisii*, making these two species an excellent platform for comparative developmental genetics studies of two closely related species with dramatic phenotypic divergence.

## METHODS

### Plant materials

The *Mimulus lewisii* Pursh inbred line LF10 (wild-type) and the mapping line SL9 were described in Yuan et al. (2013a, b). Seeds of wild *M. verbenaceus* were collected from the Oak Creek Canyon (Sedona, AZ) and the inbred line MvBL was generated by single seed descent for >10 generations. Ethyl methanesulfonate (EMS) mutants were generated using LF10 and MvBL for *M. lewisii* and *M. verbenaceus*, respectively, following Owen and Bradshaw (2011).

### Carotenoid concentration

Carotenoid pigments were extracted from the nectar guides of fresh *M. lewisii* flowers as described in Sagawa et al. (2016). Carotenoid concentration was estimated by absorbance measurement at 440 nm and normalized to 100 mg tissue. For *M. verbenaceus*, we simultaneously extracted and separated anthocyanins and carotenoids. This was accomplished by grinding two petal lobes in 200 μL methanol, which dissolves both carotenoids and anthocyanins. After 2 minutes of centrifugation at 13,000 rpm, 150 μL of the clear pigment extract were transferred to a new tube and then thoroughly mixed with 150 μL of water and 150 μL of dicholormethanol. The pigments were separated by centrifugation (13,000 rpm for 2 minutes): water-soluble anthocyanins were suspended in the aqueous phase, while hydrophobic carotenoids remained in the non-aqueous phase (Figure 7A and E).

### Bulk segregant analysis

Genetic mapping of *RCP2* followed the protocol laid out in Yuan et al. (2013b). In short, we crossed *rcp2-1* (which was produced in the LF10 background) with the mapping line, SL9, and selfed an F1 individual to produce an F2 population. We extracted DNA from 120 F2 individuals displaying the mutant phenotype and pooled the samples for deep sequencing on an Illumina HiSeq 2500 platform. We mapped the ∼196 million reads (Bioproject: PRJNA326848) to the SL9 genome using CLC Genomics Workbench 7.0 (Qiagen) and then scanned for regions enriched in homozygous single nucleotide polymorphisms (SNPs). This allowed us to identify a 70-kb candidate region.

### Quantitative RT-PCR

We extracted RNA and synthesized cDNA according to Yuan et al. (2013a). The relative transcript level of the corolla-expressed paralogs of the carotenoid biosynthetic genes (see Sagawa et al., 2016), as well as the *RCP1* and *RCP2* genes, was assessed by quantitative reverse transcriptase PCR (qRT-PCR) (primers are listed in Supplemental Table 1). qRT-PCR was performed using Power SYBR Green PCR Master Mix (Applied Biosystems) on a CFX96 Touch Real-Time PCR Detection System (Bio-Rad). Samples were amplified for 40 cycles of 95° for 15 s and 60° for 30 s. Amplification efficiencies for each primer pair were determined using critical threshold values obtained from a dilution series (1:4, 1:8, 1:16, 1:32) of pooled cDNAs. *MlUBC* was used as the reference gene as described in Yuan et al. (2013a). Three biological replicates were used for all qRT-PCR experiments. Relative expression of each target gene compared to the reference gene was calculated using the formula (*E*_ref_)^CP(ref)^ / (*E*_target_)^CP(target)^.

### RNAi plasmid construction and plant transformation

We built an RNAi construct by cloning a 408-bp fragment of exon 23 of the *M. lewisii RCP2* gene into the pFGC5941 vector (Kerschen et al., 2004) in both the sense and antisense directions (Primers are listed in Supplemental Table 2). This fragment, and every 12-bp block within it, matched only a single region of the *M. lewisii* (100% identity) and *M. verbenaceus* genomes (95% identity), indicating target specificity. The plasmid was verified by sequencing and then transformed into *Agrobacterium tumefaciens* (strain GV3101), before being transformed into wild-type LF10 and MvBL plants by vacuum infiltration following the protocol described in Yuan et al. (2013a).

### Protein subcellular localization

To determine the subcellular localization of RCP2, the full-length *RCP2* coding DNA sequence (CDS) was cloned into the Gateway vector pEarleyGate 104 (Earley *et al*., 2006), which drives the expression of the transgene by the CaMV 35S promoter and fuses the YFP CDS in frame with the 5’-end of the target gene CDS (Primers see Supplemental Table 2). The *35S:YFP-RCP2* plasmid was sequence verified before being transformed into *Agrobacterium tumefaciens* (strain GV3101). The *35S:RCP1-YFP* plasmid was generated previously (Ding and Yuan, 2016), and was used as a positive control. For transient protein expression, *Agrobacterium* solutions containing the *35S:RCP1-YFP* or *35S:YFP-RCP2* plasmid were injected to the abaxial side of *Nicotiana benthamiana* leaves, following Ding and Yuan (2016). Fluorescence images were acquired using a Nikon A1R confocal laser scanning microscope equipped with a 60X water immersion objective.

### Yeast two-hybrid assay

Yeast two-hybrid constructs were built using the Matchmaker Gold Yeast Two-Hybrid System (Clontech). The full-length *RCP2* CDS was recombined into the pGBKT7-BD bait vector using the In-Fusion Cloning Kit (Clontech) and then transformed into the Y2H Gold yeast strain by PEG transformation, according to manufacturer’s instructions. The *RCP1* CDS was recombined *in vivo* into the pGADT7-AD prey plasmid in the Y187 yeast strain (primers listed in Supplemental Table 2). Both plasmids were brought together in individual yeast cells by mating between the two yeast strains and screened on DDO, QDO, and QDO/X/A plates to test for protein-protein interactions.

### Transmission electron microscopy

Pieces of nectar guide tissue of *M. lewisii* or petal lobe tissue of *M. verbenaceus* were pre-fixed in 2.5% glutaraldehyde and 2.0% paraformaldehyde with 0.05 M PIPES buffer. The samples were post-fixed with 1% osmium tetroxide and 0.8 % K_3_Fe(CN)_6_ and then dehydrated in ethanol. Samples were embedded in Spurr’s resin and sectioned tangentially. Sections were counterstained with 2% aqueous uranyl acetate and 2.5% Sato’s lead citrate. The sections were examined and photographed on the FEI Tecnai 12 G2 Spirit BioTWIN transmission electron microscope at UConn’s Bioscience Electron Microscopy Laboratory.

## ACKNOWLEDGEMENTS

We thank Clinton Morse, Matt Opel, and Adam Histen for plant care in the UConn EEB Research Greenhouses, thank Maritza Abril and Xuanhao Sun at the UConn Bioscience Electron Microscopy Laboratory for assistance in the TEM experiments. This work was supported by the University of Connecticut start-up funds and an NSF grant (IOS-1558083) to Y-W.Y.

